# Regions with intermediate DNA methylation are epigenomic hotspots governing behavioral adaptation to changing environments

**DOI:** 10.1101/2022.12.11.519758

**Authors:** Elizabeth Brindley, Faten Taki, Shannon C. Odell, Madelyn R Baker, Judit Gal Toth, Miklos Toth

**Author notes:** Corresponding author Miklos Toth.

## Abstract

Epigenetic and associated gene expression changes in the brain drive animals’ ability to adapt to changing environments. However, epigenetic attributes of environmental adaptation are unknown. Here we show that exercise, unpredictable stress, and environmental enrichment, conditions that elicit adaptive changes in synaptic plasticity and spatial learning, result in CpG methylation changes in regions that exist in both methylated and unmethylated states (i.e., epigenetically bistable) in hippocampal granule cells. Sustained exposures altered both the epiallelic proportions at these regions and neuronal and behavioral adaptation, indicating their adaptive nature. These malleable regions were enriched in exons and enhancer-associated chromatin marks. Their locations were mostly unique to specific environments but converged on similar synaptic genes. Lastly, manipulating DNA methylation altered epiallelic proportions at bistable regions in granule cells and phenocopied adaptive behavior. We propose that shifts in epiallelic proportions at evolutionarily conserved bistable regions, via gene expression changes, contribute to hippocampal plasticity and behavioral adaptation to changing environments.

## Introduction

There is an increasing body of evidence demonstrating that behavioral adaptation of an individual to a changing environment is driven by epigenetic and associated gene expression changes in the brain^1,2^. We hypothesized that adaptive responses occur through genomic regions with environmentally induced epigenetic malleability. As environmental challenges have been part of the species’ existence throughout their evolution, we reasoned that mechanisms producing and maintaining these malleable regions have evolved during natural selection and would therefore be enriched in conserved regions of the genome. Indeed, it has been shown in model organisms that initial rapid transcriptional plasticity to an evolutionarily novel environment is superseded by evolutionary/genetic adaptation that allows phenotypic adaptation in the now familiar environment^3,4^.

To identify environmentally malleable epigenomic regions, we exposed mice to environments similar to their ancestral (familiar) environment. Voluntary exercise, via free access to a running wheel, is a naturalistic behavior, as mice in the wild use it without any extrinsic reward^5^. Chronic unpredictable stress^6^, including exposure to a predator odor, temperature fluctuations, and social isolation, is also common in nature. Finally, environmental enrichment mimics the complexity of the physical and social environment of natural environments. Although studies reported running-, stress-, and environmental enrichment-associated epigenetic changes at specific genes and genome wide^7-10^, they revealed no general principles that underlie malleability in the epigenome to the environment.

The identification of epigenomic regions malleable by the environment could be complicated by the inherent differences between environmental challenges and how/where in the brain they are processed, especially between those with a hedonic (running) and aversive (stress) nature. However, all challenges have contextual and spatial components in common, which are processed by the dorsal hippocampus^11^. Indeed, running, chronic stress, and environmental enrichment all induce adaptive changes in hippocampal neurons and affect spatial/contextual memory^12-14^. We selected dorsal dentate gyrus (dDG) granule cells (DGCs) to study, as they are the first hippocampal neurons to receive contextual and spatial information from the environment^11^. Further, due to their inherently low excitability, DGCs gate the flow of spatial and contextual information from the entorhinal cortex into the hippocampus^15,16^. Of the various epigenetic modifications, we focused on CpG methylation because of its relative stability and binary nature that allows the conversion of methylation differences to proportions of cells switching methylation state. By investigating methylation changes and their genomic location in response to different environments, we identified thousands of environmentally induced differentially methylated regions (env-DMRs) and describe the criteria that define environmentally induced regional epigenetic malleability in the hippocampus.

## Results

### Changing environment alters methylation at thousands of short genomic regions

We verified that four weeks of voluntary exercise, via free access to a running wheel^17^ (**Fig. 1a**), induced hippocampal neuronal and behavioral plasticity. As reported earlier, running (∼6 min bouts for a total time of ∼3h during the night and ∼30 min during the day) produced structural changes^18,19^, specifically an increase in dendritic spine density in DGCs (**Fig. S1a**), and enhanced spatial memory in the Morris water maze (MWM) of male C57BL6 (“run”) mice^20,21^ (**Fig. S2a**). Next, from an independent cohort of run and control mice we microdissected the granule zone from dDG slices, which due to the laminar structure of the DG is comprised of the cell bodies of DGCs with minimal contamination from nuclei of other cell types. Following DNA isolation, we used enhanced reduced representational bisulfite sequencing (eRRBS)^22^, as it is a cost-effective method to achieve the sufficiently deep sequencing needed to interrogate environmentally induced changes in cytosine methylation (**Fig. 1b**). We recovered 1.5-2 million CpG sites with ≥10x coverage across samples (average of 50.3x) and identified sites with at least 15% methylation difference (q>0.01) between run and control animals. The threshold of 15% has been shown to be optimal for balancing sensitivity and specificity in differential methylation detection across RRBS analyses methods^23^ and has been widely used in DNA methylation studies^24,25^. Nevertheless, we further increased stringency in differential methylation detection by applying the 15% threshold to 2 or more clustered CpGs (within 1kb) because multiple CpG sites more reliably predict true and reproducible changes and increase the probability of functional relevance^26^. This approach identified 3,199 differentially methylated regions between run and control DGCs, hereby referred to as run-DMRs. Run-DMRs were on average 164.2 bp long and contained an average of 2.50 differentially methylated CpG sites, representing 0.65% of all CpG sites covered at ≥10x (**Fig. 1c, Table S1** for links to custom tracks in the Integrative Genomics Viewer [IGV]).

**Figure. 1.**
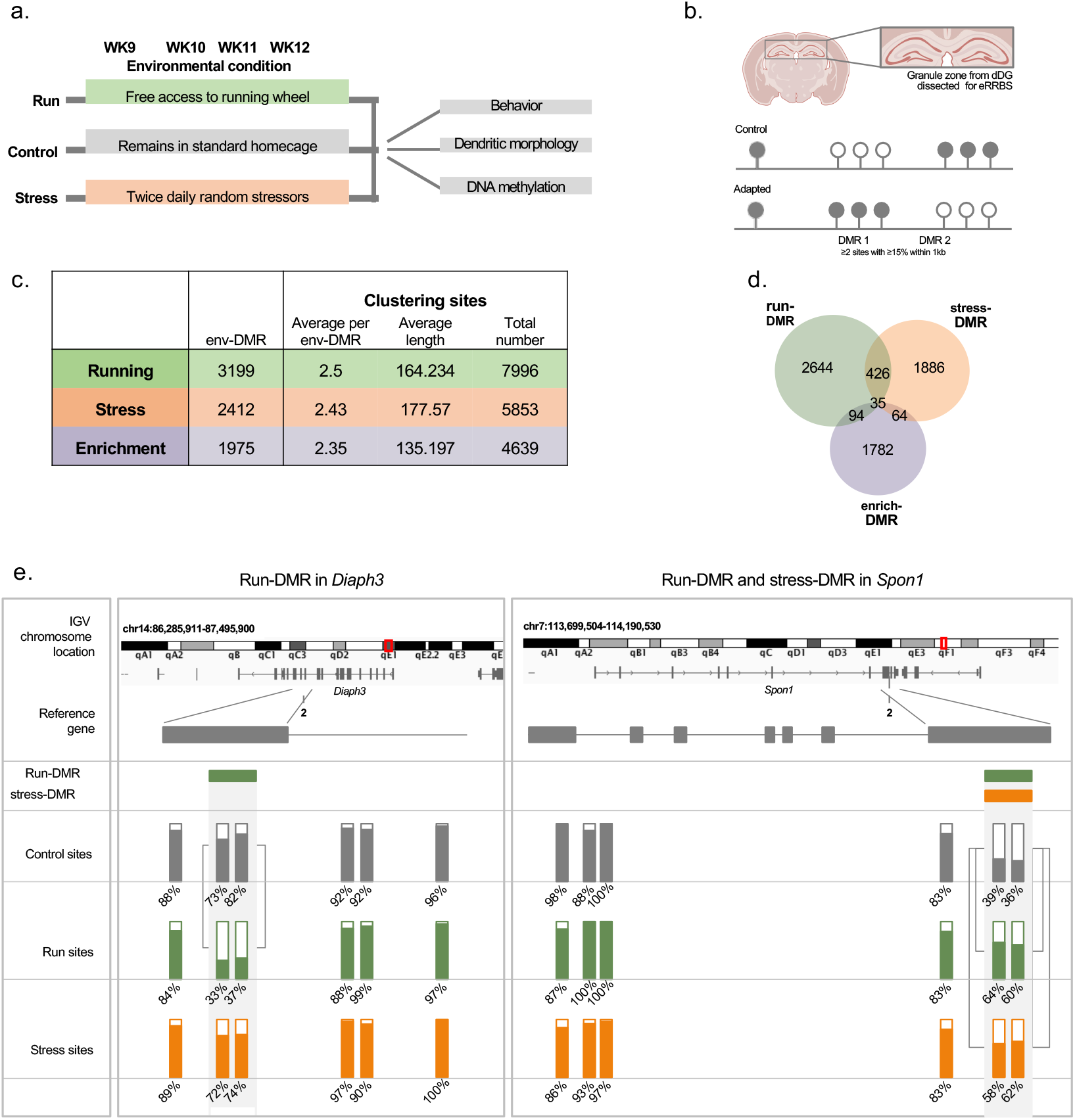
Epigenetic adaptation in the dDG to daily running and chronic stress. **a**, Experimental timeline of daily voluntary exercise via free access to a running wheel and chronic unpredictable stress and downstream experiments. **b**, Isolation of DGC cell bodies for eRRBS and schematic representation of epiallelic switching in a hypermethylated and a hypomethylated region following an environmental challenge. **c**, Summary of changes in DNA methylation induced by sustained environmental exposure. **d**, Most env-DMRs are unique to an environment as shown by the modest overlap (at least 1 nucleotide) between regions. **e**, IGV browser view and schematic of a hypomethylated run-DMR with intermediate methylation in control and run samples, a non-DMR cluster of highly methylated CpG sites with no methylation change by the environment, and a hypermethylated dual run/stress DMR with intermediate methylation in control, run and stress.

Next, we tested if an aversive environment targets epigenetic regions similar or different to those altered by the hedonic experience of exercise. The chronic unpredictable stress paradigm consisted of four weeks of twice daily exposures to a multitude of stressors in a pseudorandom order (**Fig. 1a**, Materials and Methods). We validated the effects of stress on DG neuroplasticity and behavior by demonstrating a reduction in dendritic branch points in the DG (**Fig. S1b**), and a failure to recall the target quadrant even after extended training in the MWM (**Fig. S2b**), as expected^14,27,28^. Stress can also be considered adaptive because reduced dendritic complexity and subsequent spatial memory impairment have been proposed to protect neurons against the stress-induced excess in glutamate release^14,27^. Such a mechanism may be of particular importance in the DG due to its inherent sensitivity to hyperexcitability and seizures^29^. Using eRRBS as described above, we identified 2,412 stress-DMRs, which had an average length of 177.57 bp and average of 2.43 CpG sites, representing 0.47% of all sites covered at ≥10x (**Fig. 1c, Table S1** for links to custom tracks in the Integrative Genomics Viewer [IGV]).

Environmental enrichment, like voluntary exercise, promotes neuronal plasticity in the hippocampus and enhances spatial memory and cognitive performance^13,30^. Using a publicly available RRBS dataset from the DG of C57BL6 mice^8^, we again found methylation changes clustering to thousands of short genomic regions (1,975 enrich-DMRs; **Fig. 1c**). Interestingly, the genomic overlap (at least 1 nucleotide) between run-, stress-, and enrich-DMRs was relatively modest (5-19%), but significant (hypergeometric testing; run vs. stress, run vs. enrich, stress vs. enrich, all p<0.0001), indicating that different environments target mostly different sets of environmentally malleable regions (**Fig. 1d**, see also **Fig 1e** for representative DMRs).

### Intermediate methylation is a core feature of environmental malleability

Intriguingly, the population-level methylation of the majority of run-DMR CpG sites was in the intermediate range (i.e., 15-85%) in control DGCs and remained in the intermediate range following running-induced gain or loss of methylation (**Fig. 2a**). Intermediate methylation of run-DMR CpGs contrasted the bimodal methylation, i.e., either methylated (>85%) or unmethylated (<15%), of CpGs outside of DMRs (**Fig. 2d**). Approximately half of the CpG sites gained methylation across the population of DGCs (average gain of 27.19%, with 95% of the sites gaining between 15.0% and 45.762%) while the other half lost methylation (average loss of 27.25%, with 95% of the sites losing between 15.0% and 44.402%; **Fig. 2b, c, Table S2**). CpGs with methylation at the lower end of the range tended to gain, while those in the higher methylation range lost methylation during running (**Fig. S3a**). Overall, although regions with intermediate methylation have previously been described in different tissues^31,32^, here we found that intermediate methylation has a functional correlate, i.e., environmental malleability, in DGCs.

**Figure 2.**
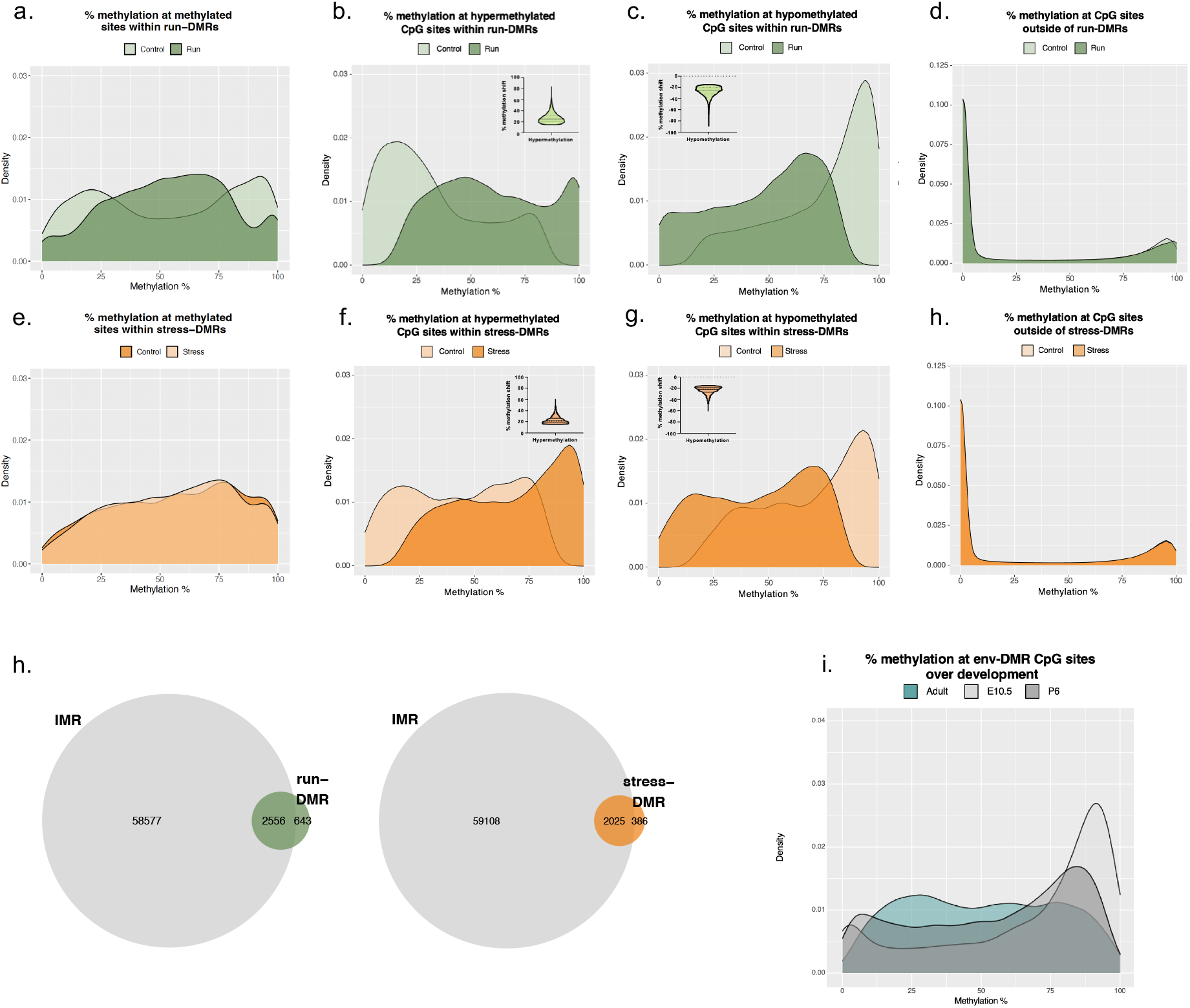
Intermediate methylation is a core feature of environmental malleability. **a-c**, Intermediate methylation of run-DMR CpGs in control and run mice and their shifts to hyper- and hypomethylation by daily running. Insets show the extent of running-induced methylation shifts with mean and s.e.m indicated. **d**, Methylation distribution of CpG sites outside of run-DMRs exhibits a bimodal distribution pattern. **e-h**, Same as “a-d” except the environmental exposure is chronic stress. **i**, Extensive overlap of at least 1 nucleotide between env-DMRs and IMRs. **j**, Intermediate methylation at env-DMR CpGs is established gradually during development. The largely bimodal methylation distribution in hippocampal progenitors (E10.5) begins to transition to intermediate methylation in young DGCs (P6), which is completed in adult mature DGCs.

Similar to run-DMRs, stress-DMRs were intermediately methylated in controls and following stress, with half of the CpG sites gaining (average gain of 23.13%, with 95% of the sites gaining between 15.0% and 36.655%), while the other half losing methylation (average loss of 23.48%, with 95% of the sites losing between 15.0% and 37.593%) (**Fig. 2e-g** and **Fig. S3b**). Further, CpG methylation outside of stress-DMRs persisted in a fully methylated or fully unmethylated state (**Fig. 2h**).

Finally, environmental enrichment also elicited methylation changes preferentially at intermediately methylated genomic regions, i.e., enrich-DMRs (**Fig. S4a-c**), and CpGs outside of DMRs were fully methylated and unmethylated (**Fig. S4d**). Taken together, these data indicate that environments with both positive (run, enrichment) and negative (stress) valence preferentially alter CpG methylation at regions that have an intermediately methylated state.

Intermediate methylation at env-DMR CpGs is due to the existence of both the unmethylated and methylated epialleles, i.e., epigenetic bistability, in the DG cell population. As the methylation state of neighboring CpGs within intermediately methylated regions is known to be similar^33,34^, bistability may be extrapolated to env-DMRs. It follows that environmental exposure shifts the native epiallelic proportions of env-DMRs and thus reprograms the epigenetic heterogeneity of the DG.

Since env-DMRs are typically intermediately methylated, we sought to computationally determine the possible repertoire of epigenetically malleable regions in naïve adult DGCs by identifying regions with at least 2 clustered CpG sites (within 1 kb), each with methylation levels in the intermediate range (between 15% and 85%). We identified ∼60,000 intermediately methylated regions (hereby referred to as IMRs) that were on average 272.93 bp long and contained an average of 4.24 CpG sites. The smaller size of env-DMRs (**Fig. 1d**) suggests that they may represent domains within IMRs that are malleable by the environmental conditions used in our experiments. As expected, the majority of run- and stress-DMRs regionally overlapped with adult DGC IMRs (**Fig. 2i**). Run- and stress-DMRs together represented 7.4% of adult DGC IMRs (9.9% when enrich-DMRs were included). IMRs, as a large set of intermediately methylated sequences, may contain additional malleable sequences poised to respond to a wide variety of environmental challenges, beyond the exposures used in our experiments.

### Short-term exposure to environment is not sufficient to alter DNA methylation

Next, we asked if short-term exposure to the above environments was sufficient to elicit run-DMRs in DGCs. We found that 24 h of wheel running produced over 10 times fewer differentially methylated regions, indicating that sustained running is required to produce DMRs (**Fig. S5a**). However, running-induced CpG methylation changes were not permanent as we detected very few DMRs in long-term Run animals (4 weeks) after an additional two months in standard cages (**Fig. S5a**).

We also tested the effect of short-term stress by exposing animals to a series of five consecutive foot shocks, using the contextual fear conditioning model. Animals were exposed to stress after a 2 min habituation period with 20 sec intervals between shocks. Foot shock control, i.e., immediate shock (IS), animals were exposed to shocks with 1 sec intervals, which is insufficient for encoding and thus forming context-shock association. As expected, context fear conditioned animals, but not IS animals, exhibited freezing in the same context one day, one week, and one month later (**Fig. S5b**). Despite this memory, we detected very few or no differentially methylated regions in DGCs one day, one week, or one month after conditioning. This indicates that although a single stress session produces long-lasting memory, it does not elicit significant methylation changes in DGCs (**Fig. S5a**). Of note, others, using controls other than those exposed to immediate shock in the conditioning context, reported methylation changes 24h after fear conditioning, possibly due to effects not related to stress-context association^35^ or to methodical differences such as the definition of DMRs. Overall, these data indicate that substantial methylation changes at clustered sites, as defined in our experiments, in DGCs require sustained exposure to an environmental challenge.

### Methylation bistability of env-DMRs and their environmentally induced methylation shifts are associated with mature neurons in the DG

Intermediate methylation at run-, stress-, and enrich-DMRs was unrelated to genomic imprinting because of its wider methylation range and different distribution across the genome. Further, intermediate methylation was not due to cellular heterogeneity as DNA samples were collected from granule cell bodies from the granule cell zone of the DG that has few other cell types and certainly much less than the 15% required to generate bistability in the population of DGCs.

However, a subset of DGCs in the adult DG are at different maturational stages, as new cells are continuously produced from late prenatal life throughout adulthood^36^. Adult-born DGCs show an initial period of hyperexcitability (up to ∼3 weeks), followed by synaptic maturation for an extended period of time (up to ∼8 weeks), before they became fully mature and functionally indistinguishable from developmentally-born DGCs that constitute the majority of the adult DG^37-39^. It is unlikely that young immature neurons are associated with our observed epigenetic bistability at env-DMRs in naïve animals or contribute to the shift in methylation following environmental exposure for several reasons. Firstly, ratios between the two epiallelic states are broad across DMR CpGs in naïve DG (15-50% for the minor epiallele per definition), and environmentally induced methylation shifts are variable from 15% up to 46% (**Fig. 2b, c, f, g**). This indicates the involvement of larger and numerically more variable cell populations than the 12-13% fraction of adult-born neurons in young adult mice, estimated by extrapolating rat neurogenesis data^40^ to mouse and adjusting for differences in survival rate in the two species^41^. Secondly, at postnatal day 6, when the DG consists of perinatally-born immature neurons (< 3-week-old), env-DMR CpGs showed a bimodal methylation pattern that was transitioning to intermediate methylation, while a truly intermediate methylation profile was evident in adult (12-week-old) mice (**Fig. 2j**). Consistent with the emergence of intermediate methylation from P6, E10.5 hippocampal progenitors showed a bimodal methylation profile at env-DMR CpGs (**Fig. 2j**). These data suggest that bistability at env-DMRs is established during maturation of DGCs and that young neurons, because of their mostly monostable methylation (and relatively small proportion, see above), may not contribute to the intermediate methylation pattern in the DG. Thirdly, while running is known to increase adult neurogenesis, stress decreases the generation of adult-born neurons, thus affecting the proportion of adult-born neurons in the DG. However, methylation shifts at run- and stress-DMRs were comparable, which again is inconsistent with the contribution of adult neurogenesis to environmentally induced epiallelic switching. Finally, we identified intermediately methylated regions in brain regions in which no adult neurogenesis is observed, such as pyramidal neurons in the CA1 and CA3, as we reported previously^33^. Overall, these data are most compatible with intermediate methylation at env-DMRs and shifts in environmentally induced methylation to be associated with the large pool of mature neurons in the adult DG. However, the size of the pool of mature neurons involved in methylation shifts is highly variable across env-DMRs.

### Methylation bistability and environmental malleability are associated with specific genomic and epigenomic features

The distribution of CpG methylation is not uniform across the genome. Promoters at active genes are typically unmethylated, while gene bodies and intergenic regions are typically methylated. Since env-DMRs are in both the methylated and unmethylated state in a population of cells, we sought to determine their genomic distribution to gain insights into their functional relevance. Run- and stress-DMRs were enriched in gene bodies, particularly in coding exons and last exon/3’UTRs (**Fig. 3a**), illustrated by representative exonic DMRs in run and stress animals in **Fig. 3c**.

**Figure 3.**
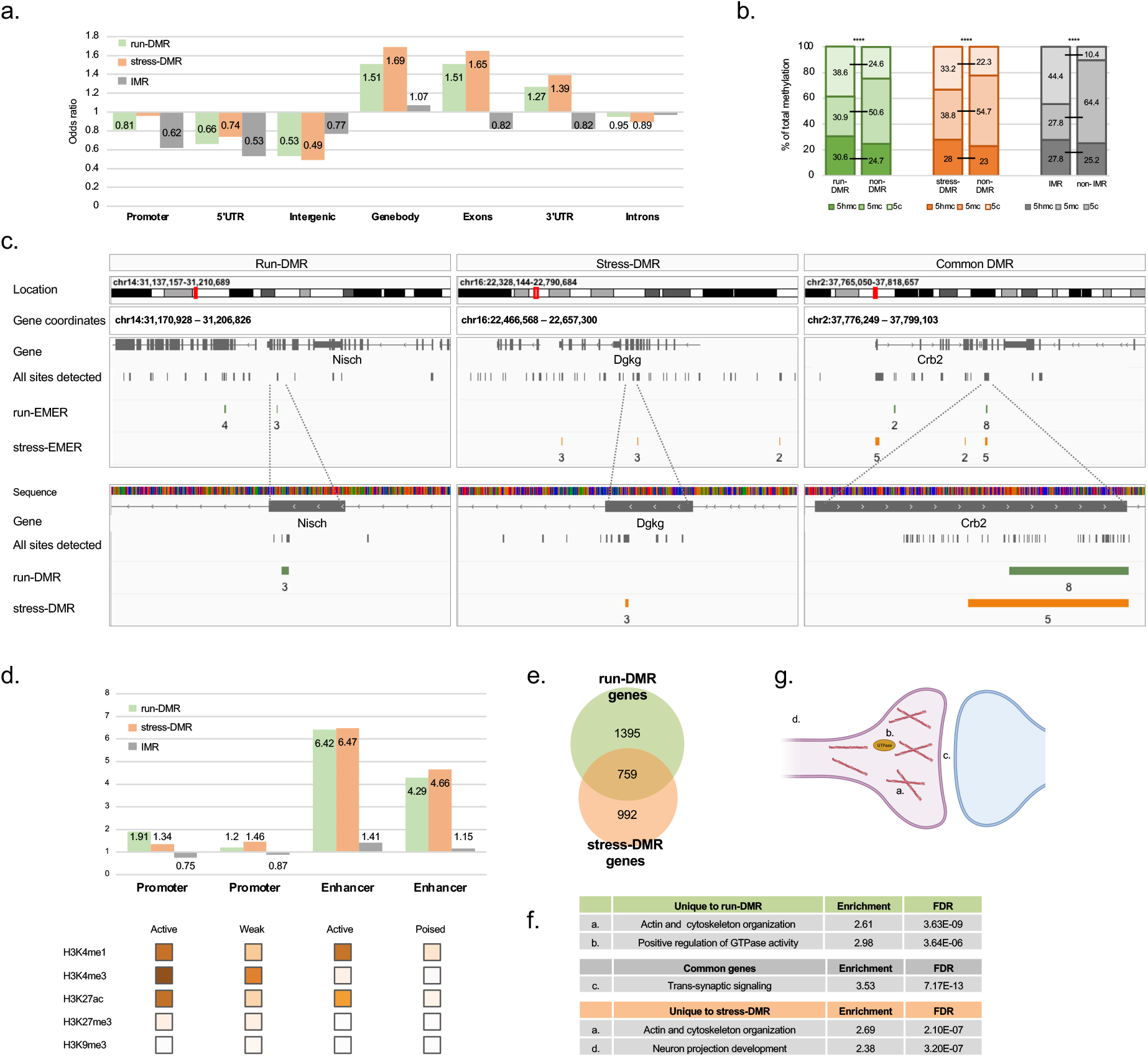
Genetic and epigenetic features associated with environmental malleability. **a**, Exonic and 3’UTR enrichment of env-DMRs/IMRs, based on odds ratio and chi square testing. All depicted values are significantly enriched/depleted (p=0.004-2.2E-16). **b**, Proportion of 5hmCs, relative to total methylation (5mC+5hmC) is higher at env-DMRs and IMRs compared to sequences outside of these regions, as determined by parallel bisulfite sequencing (BS) and oxBS. Tukey’s multiple comparisons tests. Adjusted *p*-value <0.001 in all comparisons. **c**, Representative image of exonic env-DMRs. IGV browser view and schematic of *Nisch* (containing run-DMR), *Dgkg* (stress-DMR), and *Crb2* (overlapping run- and stress-DMR). **d**, Env-DMRs are enriched in histone modifications characteristic for active and poised enhancers, based on chromHMM^46^. The heatmap, adapted from Zhu *et al*., shows the emission probability of relevant histone marks identified by chromHMM^46^. **e-f**, Venn diagram indicates substantial overlap between run- and stress-DMR containing genes, but unique genes for run and stress also define identical or similar biological processes. Only the most significant biological processes, as determined by GO analysis, are shown. **g**, Biological processes in “**f**” all map to the postsynaptic/dendritic compartment of neurons.

In contrast to their exonic enrichment, run- and stress-DMRs were depleted in promoters and 5’ UTRs suggesting that they are not associated with transcriptional initiation. Further, run- and stress-DMRs were depleted in intergenic sequences. Conversely, IMRs were present at near expected frequency in gene bodies/exons, but similar to run- and stress-DMRs, were depleted in upstream regulatory regions. Overall, run- and stress-DMRs may represent a subset of IMRs that are primarily exonic and conserved through mammalian evolution.

Exons and 3’UTRs are reportedly enriched in 5-hydroxymethylcytosines (5hmC)^43^, an intermediate produced from 5mC as it is converted to cytosine by ten-eleven translocation (TET) enzymes^44^. Therefore, we assayed env-DMRs, IMRs, and sequences outside of these regions for both 5mC and 5hmC. DMR/IMR sequences had a lower total methylation level (5mC+5hmC) than outside regions, consistent with the intermediate methylation of env-DMRs/IMRs and high methylated state of the majority of the genome. However, the relative proportion of 5hmC in total methylation was higher in both run/stress-DMRs and IMRs, as compared to the rest of the genome, indicating the enrichment of 5hmC in these intermediate methylated genomic regions (**Fig. 3b**).

Next, by using publicly available Paired-Tag data for mouse DGCs^45^, we found that run- and stress-DMRs, but not IMRs, were enriched in chromatin marks characteristic for active and poised enhancers, as determined by chromHMM^46^ (**Fig. 3d**). Since enhancers are typically located in introns and intergenic regions, the enhancer specific chromatin association and exonic enrichment of env-DMRs seem contradictory. However, ∼7% of the putative enhancers have been assigned to exons, and protein-coding sequences have been reported to function as transcriptional enhancers^47-49^. The difference in chromatin associations (in addition to the difference in genomic features) provided a further distinction between env-DMRs and the larger group of IMRs and suggest that, although IMRs may contain additional env-DMRs, not all IMRs are expected to be environmentally malleable. Overall, these data indicate that run- and stress-DMRs tend to be located in exons and may function as enhancers. Since coding exons are conserved due to the evolutionary constraint of protein coding regions, we assessed the evolutionary conservation of run- and stress-DMRs relative to control eRRBS regions of similar size. Although control sequences (non-DMR/IMR regions with clustered CpGs) showed a moderate conservation (PhastCons scores^42^ of 0.2297), due to their enrichment in relatively CpG dense and conserved regions such as promoters, run- and stress-DMRs had a significantly higher conservation score (0.3267 and 0.3257, respectively; both *P*<0.0001) suggesting the evolutionary conservation of env-DMRs across 56 mammalian species.

### Run- and stress-DMRs are both associated with synaptic genes

Run- and stress-DMRs were associated with 2,154 and 1,751 genes, respectively (**Fig. 3e**), indicating that a large number of genes contain embedded environmentally malleable epigenetic regions. However, we found no obvious correspondence between hyper- and hypomethylated DMR genes and differentially expressed genes (DEGs) in run and stress mice (**Table S3**), probably due to multiple factors, such as the complex relationship between gene body methylation and gene expression^50,51^ and as not all differential methylation may produce detectable expression changes while non-DMR genes may show compensatory differential expression. Interestingly, despite their limited genomic overlap, 35-43% of run- and stress-DMR genes overlapped, indicating that run and stress often target the same gene, but at different locations (**Fig. 3e, Table S4** for lists of genes). Analysis of this common set of 759 genes using PANTHER Gene Ontology (GO) annotations^52^ showed enrichment in the biological process of “trans-synaptic signaling” and included genes encoding neurotransmitter receptors (glutamate, dopamine, GABA), channels, and postsynaptic density proteins (Shanks; **Fig. 3f**), consistent with the central role of structural and functional synaptic plasticity in run- and stress-induced adaptation^14,18,20,53,54^.

DMR genes unique to run and stress were also enriched in a shared biological process, namely “actin cytoskeleton organization.” This finding is consistent with dendritic structural changes in run and stress mice (**Fig. 1b, e**), as actin cytoskeleton is the principal architectural component of dendritic spines^55^ and traverses, as F actin, the lengths of dendrites^56^. Finally, run- and stress-specific functions were also found, although with lower FDR. The top biological processes for running was “positive regulation of GTPase activity” and for stress, “neuron projection development,” which can also be linked to actin/spine remodeling^57^. Therefore, although run- and stress-DMRs were largely different, they converged on the same genes or biological processes, specifically on those involved in the organization of postsynaptic and dendritic structure (**Fig. 3g**).

### env-DMRs, created in the absence of any true experience, produce behavioral change

We reasoned that regions with epigenetic bistability are more sensitive to perturbations in the DNA methylation/demethylation machinery than stably unmethylated and methylated regions in the genome and that increasing and decreasing the *de novo* methyltransferase DNMT3A may shift epiallelic proportions at bistable regions. To test this assumption, adult male mice were injected bilaterally in the dDG with either DNMT3A-expressing AAV-DJ-SYN-DNMT3a-GFP virus or control AAV-DJ-SYN-GFP virus, titrated to achieve a relatively sparse (20-30%) expression pattern^33^, similar to the fraction of cells that undergo methylation changes following an environmental challenge. As we reported earlier^33^, *Dnmt3a* and control virus injected animals exhibited GFP expression in the DG twenty days later.

Overexpression (OE) of DNMT3A resulted in a total of 6,515 differentially methylated regions or OE-DMRs in the DG (**Fig. 4a, d**). OE-DMRS had a similar size and CpG density than env-DMRs (**Figs. 4d vs. 1c**). Methylation of OE-DMR CpGs shifted from a relatively low intermediate level (<50%) to a higher level within the intermediate range (**Fig. 4a**), resembling the increase in methylation at the hypermethylated fraction of env-DMR CpGs (**Fig. 4a** vs. **Fig. 2b, f**). A small fraction (∼4%) of OE-DMR CpG sites (**Table S5**) was paradoxically hypomethylated, possibly due to an imbalance of DNMT3A and TET at certain CpG sites causing loss of methylation^58^. Notably, unmethylated sequences, such as promoters and CpG islands, were mostly spared likely because of their protection against *de novo* methylation^59^. Indeed, CpG sites outside of OE-DMRs had a bimodal distribution, supporting our prediction that OE-D3A preferentially methylates CpG sites with intermediate methylation (**Fig. 4b**). Preferential targeting of intermediate methylation by OE was further supported by the high regional overlap (81.0%, p<0.0001) of OE-DMRs with IMRs (**Fig. 4c**). OE-DMRs had a significant (p<0.0001) but less robust overlap with run- and stress-DMRs (8.7% and 6.2%), which is likely an underestimation given that run-DMRs harbor not only hypermethylated but also hypomethylated CpGs (**Fig. 4c**). Indeed, the overlap between OE-DMRs and env-DMRs was comparable to that between individual env-DMRs (**Fig. 1d**) indicating that env-like DMRs can be created in the absence of an environmental change.

**Figure 4.**
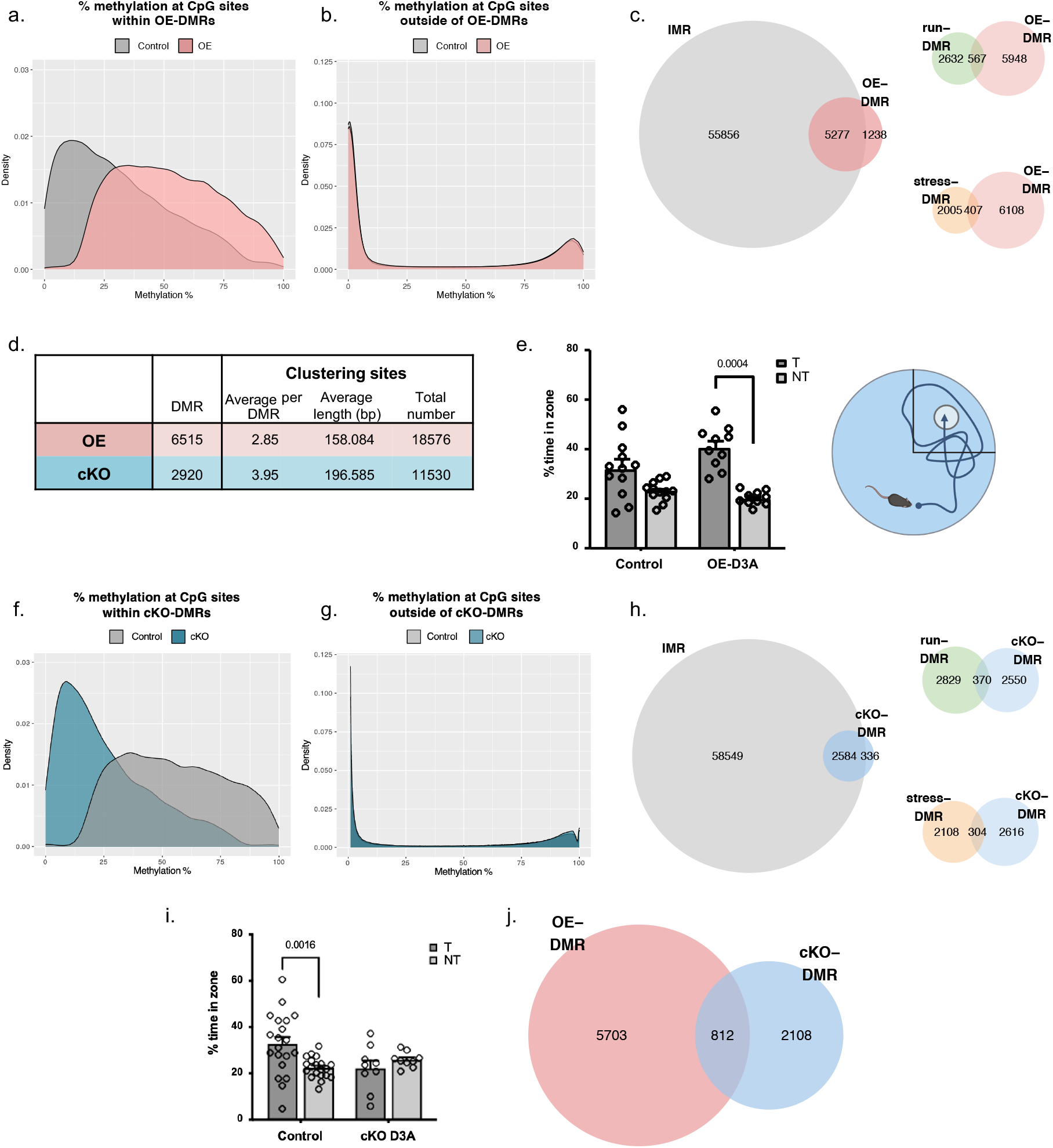
Manipulation of DNA methylation preferentially targets IMRs and alters spatial reference memory. **a**, Distribution of methylation at OE-DMR CpGs. Methylation shifts towards higher levels within the intermediate range following viral overexpression of DNMT3A in the dDG. **b**, Bimodal methylation distribution of CpGs outside of OE-DMRs. **c**, Regional overlap, based on at least 1 nucleotide, between OE-DMRs and IMRs, as well as between OE-DMRs and run- and stress-DMRs. **d**, Summary of DNA methylation changes induced by manipulation of DNMT3A expression. **e**, Spatial reference memory of OE-D3A mice in the MWM assessed by measuring time spent in target (T) quadrant in a probe trial. While a relatively limited 4-day training was not enough for control males to form memory (trend only, p=0.0659), it was sufficient for OE-D3A males to recall the platform location. Two-way ANOVA with Sidak’s multiple comparisons test; genotype x quadrant, F(1,20)=23.74, P<0.0001; n=10-12 per group, adjusted p-value indicated on graph. Error bars are mean ± s.e.m. **f**, Methylation distribution at cKO-DMR CpGs. Methylation shift towards hypomethylation within the intermediate range following cKO of *Dnmt3a* in the dDG. **g**, Bimodal methylation distribution at CpG sites outside of cKO-DMRs. **h**, Regional overlap, based on at least 1 nucleotide, between cKO-DMRs and IMRs, as well as between cKO-DMRs and run- and stress-DMRs. **i**, Impaired spatial memory of cKO-D3A mice in the probe trial of the MWM. Two-way ANOVA with Sidak’s multiple comparisons test; genotype x quadrant, F(1,54)=7.133, P=0.01; n=9 for cKO, n=20 for control group, adjusted p-value indicated on graph. Error bars are mean ± s.e.m. **j**, Overlap between cKO-DMRs and OE-DMRs.

Given that run- and stress-DMRs were associated with changes in spatial memory, we tested if non-environmental OE-DMRs have any effect on MWM memory. Following 4 days of training, OE-D3A mice had a strong and significant preference for the target zone versus non-target zones in the probe trial of MWM (in the absence of the platform), while the same training was not sufficient for control males to form memory (trend only, p=0.0659; **Fig. 4e**). Of note, performance of male mice that underwent surgery and virus delivery was reduced relative to naïve males in the MWM memory task, explaining the trend in control virus injected mice. The overall distance travelled in the MWM did not differ between OE-D3A and control mice (1,075 and 1,024cm, respectively; p=0.4574, t=0.7643, df=14), indicating that activity in the maze did not influence memory recall. Further, OE-D3A mice exhibited freezing indistinguishable from that of control (virus) mice one day after contextual fear conditioning (that requires the coordinated activity of the ventral CA1 region and basolateral amygdala^60,61^), indicating that enhanced performance in the MWM was not due to a generalized increase in memory formation (**Fig. S6a**). Overall, these data suggest that prior increase of methylation at a subset of regions with intermediate methylation by OE in the DG enhances spatial reference memory formation.

Next, we tested if neuron specific genetic inactivation of *Dnmt3a* in nestin-creERT2 mice^33,62-64^; (cKO) would reduce methylation at IMRs. Deletion of *Dnmt3a* postnatally (from ∼two weeks of age) was reported to have no effect on DNA methylation in the hippocampus^65^. However, cKO of *Dnmt3a* by a single dose of tamoxifen (TAM) at E13.5, before the emergence of IMRs during development (**Fig. 2j**), reduced methylation from an intermediate to a lower level at 2,920 cKO-DMRs in adult mice (**Fig. 4d, f**). Only a small fraction (1.2%) of cKO-DMR CpG sites were hypermethylated (**Table S5**). CpGs outside of cKO-DMRs retained their bimodal distribution indicating that cKO did not interfere with methylation at fully methylated sites (**Fig. 4g**), presumably due to the partial nature of D3A-cKO and the presence of the maintenance methyltransferase DNMT1. Selective hypomethylation at IMRs was also supported by the high regional overlap (88.5%, p<0.0001) of cKO-DMRs with IMRs (**Fig. 4h**). cKO-DMRs had a less robust but still significant regional overlap with run- and stress-DMRs (12.6% and 10.4%, p<0.0001, **Fig. 4h**), which again is likely an underestimation given that stress-DMRs contain not only hypomethylated but also hypermethylated CpGs. The cKO-DMR and IMR overlap is similar to the overlap between the individual env-DMRs (**Fig. 1d)**, indicating that env-like DMRs can be created by reducing methylation in the absence of an environmental change.

Testing spatial memory in MWM probe trials revealed that while control animals explored the area that contained the platform during training, cKO mice had no preference for this target quadrant (**Fig. 4i**). The overall distance travelled in the MWM was not different between cKO and control mice (847 and 963 cm, respectively; p=0.2784, t=1.106, df=27). Behavior of cKO mice was not affected in the contextual fear conditioning test (**Fig. S6b**), suggesting the specificity of memory impairment in the MWM. Taken together, bidirectional manipulation of DNMT3A expression resulted in increased and reduced methylation at IMRs that partially overlapped (27.5% of cKO-DMRs with OE-DMRs, p<0.0001, **Fig. 4j**) and led to opposing performance in the MWM. These studies strongly link intermediate methylated regions in DGCs to adaptive behavioral plasticity.

## Discussion

Here, we identified thousands of small genomic regions whose methylation is environmentally malleable in hippocampal DGCs. We propose that these environmentally malleable epigenetic regions allow for adaptive changes in neuronal structure and function and animal behavior in response to sustained environmental exposure, in the form of voluntary exercise and chronic stress.

The key characteristics of env-DMRs that distinguish them from the rest of the genome is their intermediate methylation, i.e., epigenetic bistability in the native state. The methylated state consists of 5mC or 5hmC, and the 5-hydroxymethylated fraction of methylation is especially prominent in env-DMRs, as well as in IMRs in general. 5hmC represents an epigenetic signal different from the typically repressive^66-68^ 5mC mark that can have implications on how the methylated state of env-DMRs may regulate gene expression^69-72^. Bistability develops during neuronal maturation during postnatal life and seems to be associated with fully mature neurons in the adult DG. Bistability of the methylation state at env-DMRs suggests a finely tuned stochastic balance between DNMT3A and TET enzymes. DMRs undergo shifts in CpG methylation in response to sustained, but not acute, environmental challenges, nonetheless DMRs remain within the intermediate range. This indicates that only a fraction of cells undergoes methylation changes at a given env-DMR. However, this fraction is highly variable across env-DMRs resulting in a complex pattern of DNA methylation states in individual neurons following environmental challenges. Our findings are most compatible with a model of environmentally induced changes in epiallelic proportions that restructure epigenomic and cellular diversity in the hippocampus promoting behavioral adaptation to changing environments.

The naïve DGC methylome contains over 60,000 regions with intermediate methylation (i.e., IMRs) that have the potential to be malleable in response to external factors. The three environmental conditions in our study identified 9.9% of IMRs exhibiting malleability. Additional environmental challenges or situations may increase the pool of epigenetically malleable IMRs. Indeed, although we included both hedonic and aversive context-related exposure, the hippocampus can also represent the social environment^2^. However, it is unlikely that all IMRs are malleable or malleable at the same degree as run- and stress-DMRs, because of their distinct differences beyond intermediate methylation. Run- and stress-DMRs tend to be concentrated in gene bodies, particularly in coding exons and last exon/3’UTRs, while IMRs are equally distributed along gene bodies. Further, run- and stress-DMRs but not IMRs are associated with enhancer-specific chromatin marks. By using these criteria as factors for environmental malleability, we predict that approximately one quarter of IMRs (including the 9.9% identified as env-DMRs) may be sensitive to environmental changes.

Based on their chromatin associations and higher 5hmC content, env-DMRs likely function as enhancers. Their exonic enrichment indicates that env-DMRs function as exonic enhancers, a group of enhancers that have a higher conservation in mammals than typical intronic and intergenic species-specific enhancers^47-49^.

Since manipulation of DNA methylation in the DG produced env-like DMRs in the absence any environmental influence and resulted in changes in spatial memory, we propose that env-DMRs and their epiallelic changes by the environment contribute to behavioral adaptation. Specifically, OE of DNMT3A increased spatial memory, similar to the adaptive enhancement of spatial memory following weeks of voluntary exercise in the running wheel. Although running produced both hypo- and hypermethylated run-DMRs, previous studies reported that increased, rather than decreased, methylation favors memory formation^64,65,73^. Therefore, we hypothesize that OE- and run-induced hypermethylation at DMRs in DGCs “primes” memory formation during subsequent MWM trials. However, the overlap between run-DMRs and OE-DMRs was relatively modest indicating that a similar behavioral output may be achieved by the hypermethylation of only partially overlapping sets of bistable regions.

Conversely, the spatial memory deficit of D3A-cKO mice may conceivably be explained by hypomethylated IMRs interfering with DNA methylation during MWM learning that otherwise is required for memory formation. Stress-induced hypomethylation of DMRs may operate similarly to impair spatial memory. Again, the relatively distinct sets of cKO- and stress-DMRs suggest that hypomethylation of only partially overlapping DMRs can lead to similar behavioral consequences. Future studies will be needed to specify the core sets of bistable regions whose methylation changes drive the behavioral changes.

In summary, env-DMRs are defined as epigenetically malleable regions responding to sustained environmental exposure by epiallelic shifts in a neuronal population. We are, for the first time, reporting the specific attributes that make genomic regions epigenetically malleable that include intermediate DNA methylation, exonic enrichment, relatively high 5hmC content, and association with enhancer-specific histone marks. We identified three sets of env-DMRs corresponding to specific environmental exposures and expect to find additional DMRs among IMRs in response to other exposures. Finally, conservation of env-DMRs and their potential role in neuronal and behavioral adaptation are consistent with similar adaptive responses of mammalian species to exercise and chronic stress and environmental challenges in general.

## Methods

### Animals

Animal experiments were performed in accordance with Weill Cornell Medical College Institutional Animal Care and Use Committee guidelines. Mice were group housed in a climate-controlled environment, 2-5 animals per cage, with a 12 h light/dark cycle (6AM-6PM). Food and water were available *ad libitum*. Adult males (between 10 and 20 weeks of age) were used for all experiments. All behavioral tests were conducted during the animals’ light phase, between 08.00 and 06.00 and all mice were habituated to the behavior room for at least 1 h prior to testing.

### Sustained voluntary exercise

8-week-old male C57BL/6J mice were obtained from The Jackson Laboratory. Following a 7-day acclimation period in standard cages, mice were placed in cage equipped with running wheels and were allowed free access for a 4-week period as previously described ^6^. Animals used the wheels, running in ∼6 min bouts for a total time of ∼3 h during the night and ∼30 min during the day, without apparent habituation throughout the 4 weeks period. Control mice remained in the standard cages for 4 weeks. A cohort of mice were euthanized following the 4-week period. A second cohort of mice were returned to standard home cages following 4 weeks of running for 2 months before being euthanized for further analysis.

### Short-term voluntary exercise

Mice were placed in cage with free access to running wheels for 24 hours (Coulbourn Instruments, Whitehall, PA).

### Chronic unpredictable stress context association

Mice were exposed to daily chronic unpredictable stress for 4 weeks based on a previously published paradigm^74^. Mice were exposed to twice daily stressors (AM and PM) from the following list; exposure to bobcat odor, cage shaking, exposure to cold (9°C for 1 h), food deprivation, forced swim, lights off, overnight cage tilt, overnight light on, overnight overcrowding, overnight strobe, restraint stress, wet bedding, exposure to white noise (3-5 h). A cohort of mice were euthanized following the 4-week period for further analysis.

### Short-term stress by foot shock

Mice allowed to habituate for 2 mins in the context before receiving a total of 5 shocks with 2 mins intervals between each^75^. The entire trial was 12 mins. Control mice received 5 shocks, with 1 sec intervals between each, immediately after entry to the context (Immediate shock or IS controls). Control mice remained in the context for 12 min.

### Overexpression of DNMT3A

Intracranial virus injection to overexpress DNMT3A in the dDG aw previously described^33^. C57BL/6 males (Taconic Biosciences) were injected bilaterally, 500 nL/side with a rate of 50 nL/min into the dDG at the following coordinates: 1.94 mm anterior-posterior, 1.0 mm medial-lateral, and 1.85 mm dorsal-ventral. The following viruses were used; DNMT3A overexpressing virus, AAV-DJ-Syn-Dnmt3a-T2a-GFP (1.2E+07 IU./mL), or control virus, AAV-DJ-Syn-GFP (6.81E+08 IU/ml). Viruses were prepared by the Stanford Vector Core.

### Genetic knock down of DNMT3A

Mice carrying floxed Dnmt3a alleles (Dnmt3a^f/f^ ; provided by Riken BioResource Center;^63^, were crossed with mice heterozygous for the tamoxifen (TAM)-inducible nestin-cre-ERT2 transgene ^62^, kindly provided by Luis Parada (University of Texas Southwestern Medical Center, Dallas, TX). Both lines were on a C57BL/6 background. Cre-negative homozygous females were bred with cre-heterozygous males. Cre-mediated *Dnmt3a* knockout was induced by TAM injection at E13.5. Pregnant dams were injected with 150-ml TAM solution (1mg), prepared by dissolving TAM (Sigma-Aldrich, St. Louis, MO) in ethanol and then in sunflower oil (Sigma-Aldrich, St. Louis, MO; 9:1) solution at 6.7 mg/ml. Gestational TAM injection interferes with females’ maternal care behavior and labor, therefore, newborn pups were delivered via C-section on E20 and cross-fostered into the nest of post-partum dams. Cre+/TAM were compared to their cre-/TAM littermate controls.

### Morris Water Maze

Morris water maze was performed as previously described^76^. Mice were trained for four days with the platform in the NW quadrant of the maze. The probe trial was run 24h after the last training trial with the platform removed. A further two days of training performed, and a second probe trial was run 24h later. Animals were tracked using Ethovision software. Time in each zone was recorded for 60s. Sidak’s or Tukey’s multiple comparisons test was performed to assess significant differences in time spent in Target versus non-target or NW and other quadrants, respectively.

### Contextual fear conditioning

Context fear conditioning was performed as previously described ^75^, with minor modifications. Fear conditions was performed in a mouse test cage (Coulbourn Instruments) inside a sound-attenuated box (Context A). Mice were habituated the chamber for 2mins before exposure to two 1-s, 0.7-mA foot shocks, delivered through the electrified floor grid each 30-s apart, each paired with a tone. Mice remained in the chamber for 1 min before being returned to their home cage. Between mice, the chamber is cleaned with 70% ethanol. 1 day, 1 week and 1 month following initial conditioning, mice were returned to Context A and freezing behavior was assessed for a 3 min period. Trials were recorded using FreezeView software (Coulbourn Instruments) and percentage time freezing was calculated. Sidak’s multiple comparisons test was performed to assess significant differences in freezing between control and fear conditioned mice.

### Dendritic complexity and Sholl analysis

Fresh brains were submerged in Golgi-Cox reagent from the FD Rapid GolgiStain Kit (Neurodigitech, San Diego, CA) solution at 25 °C in the dark for 14 days as described previously^77^. 150-μm coronal serial sections were prepared and quantitative microscopy was performed on the dDG with Microbrightfield (MBF Bioscience). Neurons were chosen by systematic random sampling. ‘Traceable’ neurons and their spines were reconstructed three dimensionally with the Neurolucida system (MBF Bioscience). A minimum of 3 cells/animal were analyzed for an n of 6 animals/group. For each neuron, 3 dendritic segments were analyzed with NeuroExplorer (Next Technologies, Madison, AL). Spine density was calculated as the number of spines per 10 µm of dendritic length. Dunnett’s multiple comparisons test was performed to assess significant differences in spine number. A two-way ANOVA, with Fishers LSD test was performed to assess differences in number of branch points in concentric circles from the soma.

### RNA extractions

Animals were perfused with 30% RNAlater (Ambion, Grand Island, NY) diluted in saline. Brains from adult mice were collected, flash frozen on dry ice and sectioned into 200 mM slices on the cryostat. The dDG was micro-dissected from the sections. Total RNA was isolated from the dDG using the RNeasy Mini Kit (Qiagen, Valencia, CA).

### Differential gene expression

Single-end, 50bp RNA sequencing was performed on Illumina HiSeq2000 and 2500 machines and aligned to the mm9 reference genome using TopHat software version 2.0.11^78^. Default parameters were used with the addition of “--no-novel-juncs” for alignment to known genes and isoforms. Genes were counted using HT-seq program^79^ with the parameter “intersection-strict”. Values for gene expression were calculated using EdgeR^80^ package in R using tagwise dispersion and default parameters. Differentially expressed genes were determined using Benjamini-Hochberg corrected *p*=0.05 threshold.

### DNA extractions

Brains from adult mice were collected, flash frozen on dry ice and sectioned into 200 mM slices on the cryostat. The dorsal dentate gyrus was micro-dissected from the sections. DNA was isolated using the QIAamp DNA Micro Kit (QIAGEN) according to manufacturer instructions with minor modifications.

### Reduced Representation Bisulfite Sequencing (RRBS)

Library preparations, sequencing and adapter trimming was performed by the Epigenetics Core at Weill Cornell Medicine as described previously^22^. A starting input of 50ng of genomic DNA, from 3-4 adult (8-12 week old) male mice was used. Single end 50 bp RRBS sequencing was performed using an Illumina 2500 according to the manufacturer’s instructions. UCSC mm10 was prepared and indexed followed by alignment and methylation calling using Bismark v22.1 using the default. MethylKit was used to perform differential methylation and statistical analyses^81^. Differentially methylation sites were defined as sites with a 15% difference in methylation between 2 groups of interest with a sliding linear model (SLIM)-corrected p-values (or q values) were greater than 0.01. Differentially methylated regions were defined as regions with two or more differentially methylated sites with 1 kb of each other^33^.

Given that the environmental enrichment dataset, unlike the run and stress datasets, was not collected in our lab and was performed with DNA from whole DG (rather than from isolated DGC cell bodies like in our study), we note these factors as potential confounds. However, methylation levels within env-DMRs and their distribution as well as the DMR overlaps were comparable across the datasets. Nevertheless, downstream analyses were performed with run-DMR and stress-DMRs only.

Genomic coordinates, exons, introns, 3’UTR, 5’UTR and TSS were downloaded from USCS genome browser based on mm10. Promoter were defined as regions ± 500 bp from the transcription start site (TSS). To evaluate enrichment or depletion of DMRs in a specific feature, we calculated the Odds Ratio as [(Number of DMR overlapping with a feature)/(Number of DMR not overlapping with a feature)]/[(Number of potential DMR overlapping with a feature)/(Number of potential DMR not overlapping with a feature)]. Odds ratios >1 and <1 indicate enrichment or depletion of DMR in a genomic feature, respectively. Statistically significant enrichments were computed using Chi-square test in R^33^.

### Reduced Representation Oxidative Bisulfite Sequencing (RRoxBS)

Bisulfite sequencing (BS) does not discriminate between 5hmc and 5mc, and the RRoxBS detects only 5mc. Therefore, 5hmc values can be calculated by subtracting 5mc levels from total methylation levels. Library preparations, sequencing and adapter trimming was performed by the Epigenetics Core at Weill Cornell Medicine as previously described^82^. A starting input of 100ng of genomic DNA used. Library preparations were performed using Ovation Ultralow Methyl-Seq DR Multiplex with TrueMethyl oxBS kit and workflow (Tecan, Redwood, CA). Single end 50 bp RRBS sequencing was performed using an Illumina 2500 according to the manufacturer’s instructions. UCSC mm10 was prepared and indexed followed by alignment and methylation calling using Bismark v22.1 using the default options^22^. Global CpG reports were used to compute percent CpG methylation and total coverage with base-pair resolution. To compute contribution of 5hmc to total methylation at a CpG sites, we considered only sites detected in BS and oxBS experiment, with a delta methylation where BS – ox BS >0.

## Data availability

DNA methylation data that support the findings of this study will be deposited to GenBank and are available from the corresponding author upon reasonable request.

## Acknowledgments

We would like to thank the following organizations for their support: The Epigenomics Core of Weill-Cornell Medicine, The Applied Bioinformatics Core of Weill Cornell Medicine, The University of Pennsylvania Vector Core, and the Stanford Vector Core. We would like to thank Grace Petkovic and Frank Nagy for their help in the chronic unpredictable stress paradigm and fear conditioning, respectively. We would also like to thank Shifra Liba Klein for initial computational analyses.

## Author Contributions

E.B., F. T., and M.T. designed research, E.B., F. T., S.C.O., and J.G.T. performed research, E.B., M.R.B, and F. T. analyzed data, E.B., M.R.B., and M.T., wrote the paper.

## Materials & Correspondence

Miklos Toth

