## Supplemental figures and tables for "Regions with intermediate DNA methylation are epigenomic hotspots governing behavioral adaptation to changing environments"

**Supplementary Figures**

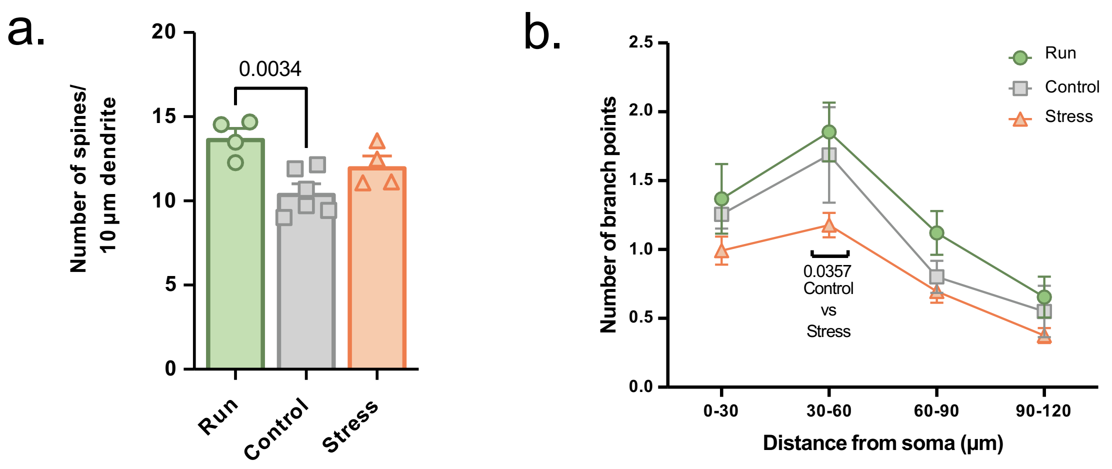


**Figure S1. Neuronal adaptation in the dDG to daily running and chronic stress. a,** Running significantly increased spine density in the dDG as shown by the number of dendritic spines per 10 µm dendrite. One-way ANOVA with Dunnett's multiple comparisons test, group x number of spines, F(2,11)=8.454, P=0.0034, n=4-6 per group, adjusted *p*-values indicated on graph. Error bars are mean ± s.e.m. **b,** Chronic unpredictable stress reduced dendritic complexity in the dDG as measured by Sholl analyses. Two-way ANOVA, with Fisher LSD test, group x distance from soma, F(2,56)=6.548, P<0.001, F(2,56), P=0.0060, N=5-6 per group, p=0.0357.

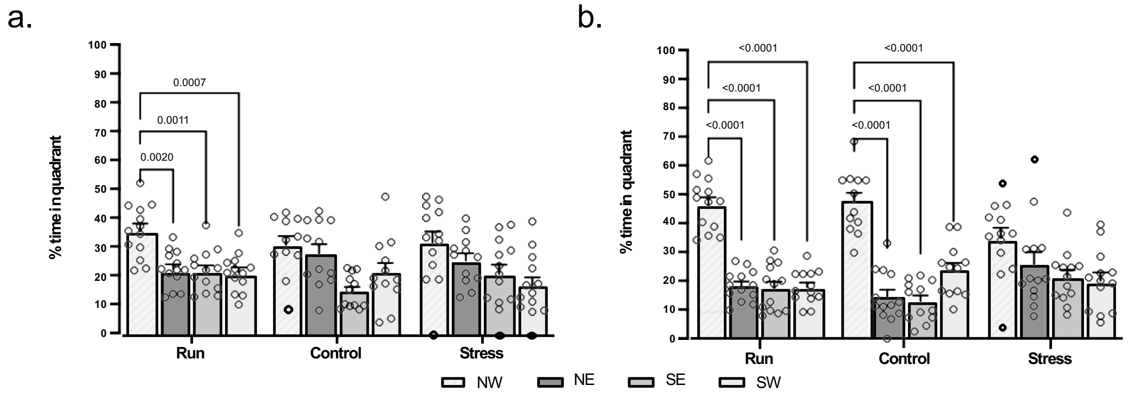


**Figure S2. Voluntary exercise and chronic stress effect spatial memory. a,** Spatial reference memory after four days of training with mice exposed to four weeks of voluntary exercise and chronic stress. Memory of mice was assessed by measuring time spent in target quadrant (NW) of the Morris Water Maze. Two-way ANOVA quadrant $F(3, 138)=15.74$, $P<0.0001$. Tukey's multiple comparisons test, adjusted p-values indicated on graph, N=12-13 per group. Error bars are mean ± s.e.m. **b**, Same as in "a", but after an additional two days of training. Two-way ANOVA quadrant $F(3, 141)=57.72$, $P<0.0001$; group x quadrant interaction $F(6, 141)=4.769$, $P=0.0002$.

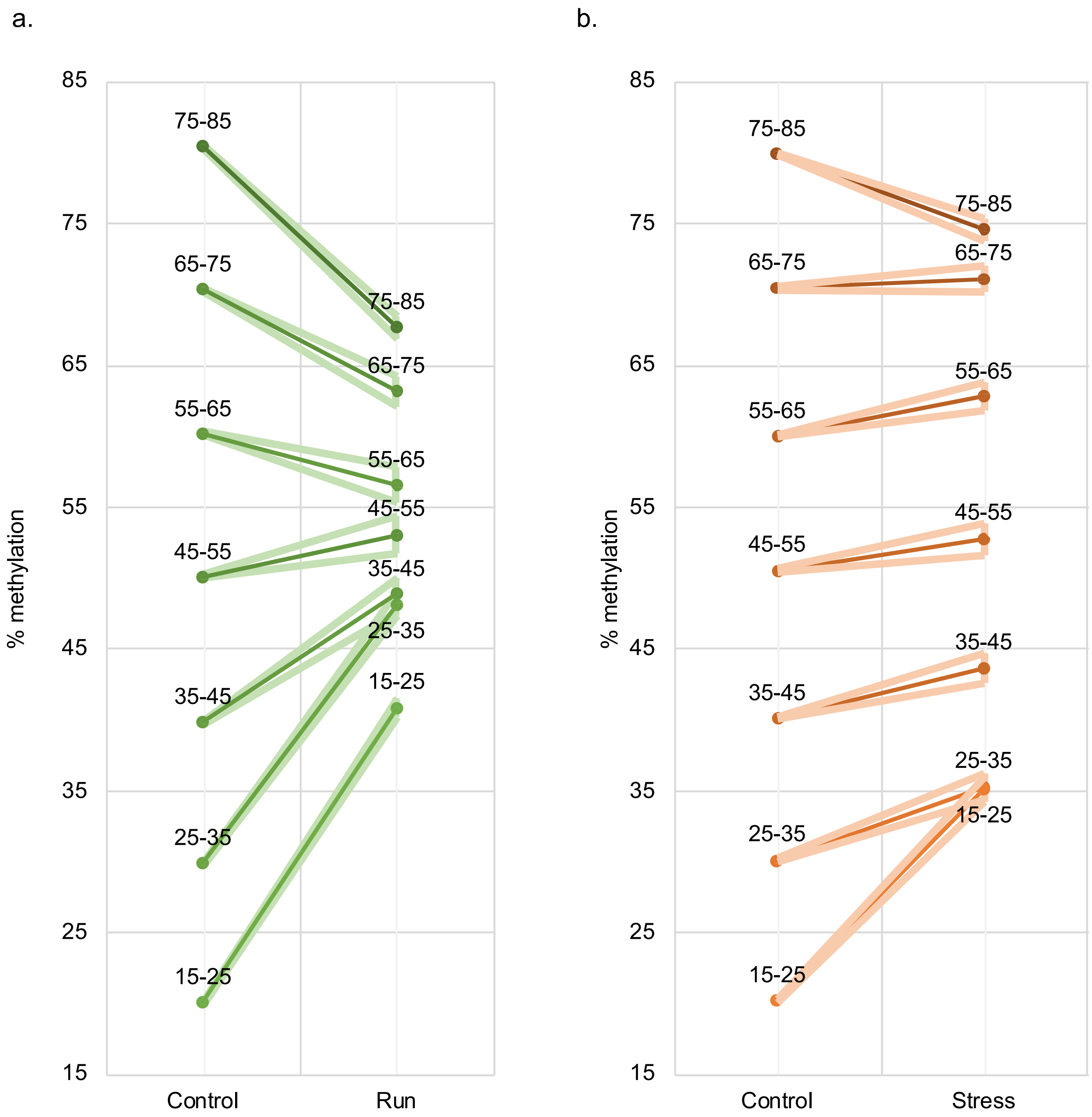


**Figure S3. Directionality of environment-induced methylation changes depends on the native methylation state of env-DMRs. a, b,** Run-/stress-DMRs with low methylation in controls tend to be hypermethylated, while those with high methylation tend to be hypomethylated following four weeks of running and chronic stress.

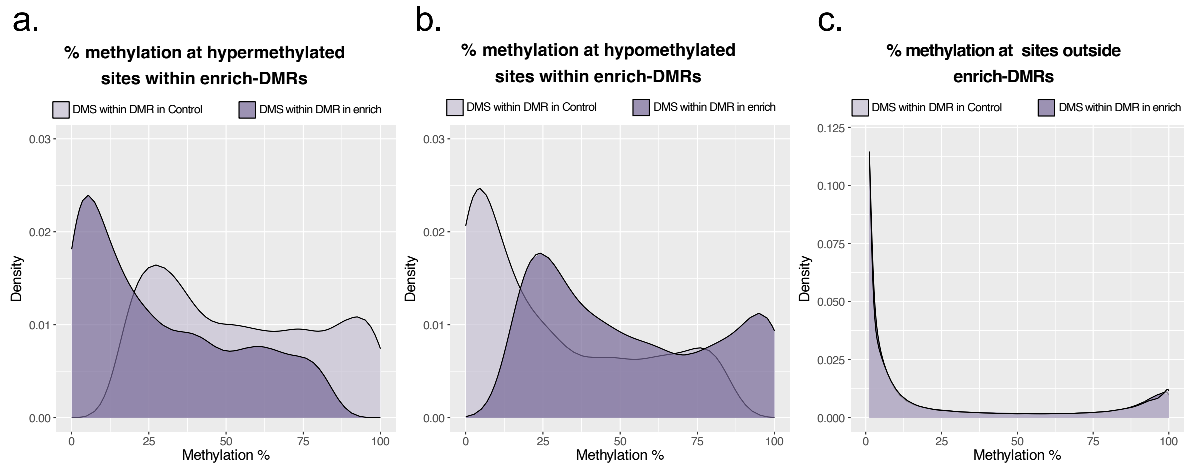


**Figure S4. Intermediately methylated regions are also malleable by environmental enrichment. a-c,** Environmental enrichment shifts methylation distribution at enrich-DMR CpGs to hyper- and hypomethylation, mostly within the intermediate range. **d,** Methylation distribution of CpG sites outside of enrich-DMRs exhibits a bimodal distribution pattern.

a.

| | Time point | DMR | Clustering sites | | |
|---|---|---|---|---|---|
| | | | Average per DMR | Average length (bp) | Total number |
| **Short-term running** | 1 day | 218 | 2.33 | 133.298 | 502 |
| **Long-term running** | 2month | 677 | 2.35 | 115.358 | 5264 |
| **Short-term stress** | 1 day | 3 | 2.25 | 17.000 | 13 |
| **Short-term stress** | 1week | 0 | / | / | 0 |
| **Short-term stress** | 1 month | 8 | 4.33 | 28.667 | 18 |

b.

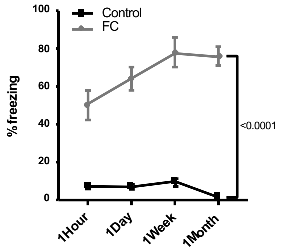


**Figure S5. DNA methylation changes at env-DMRs require long-term exposure and are reversible. a,** Effect of short-term (one day) running on DGC DNA methylation. The table also shows the reversibility of methylation changes induced by long-term running following 2 months in the home cage. Further, acute stress (one session of 5x foot shock) causes minimal DNA methylation changes measured 1 day, 1 week, and 1 month after the stress in the home cage. **b**, One session of 5x foot shock elicits strong and lasting fear memory.

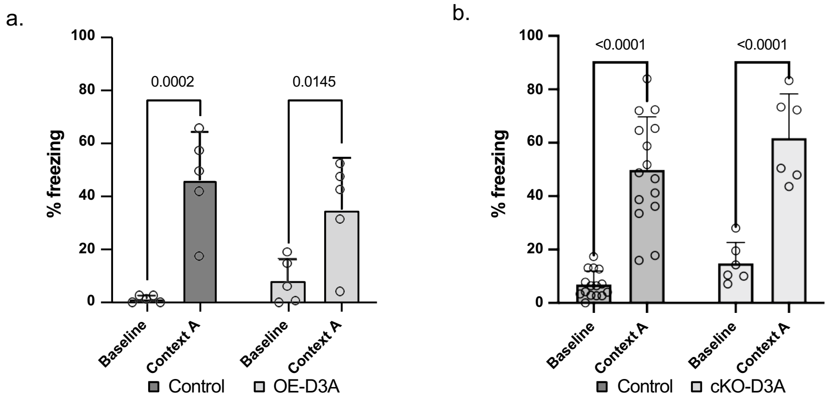


**Figure S6. Fear memory is not affected by the manipulation of DNMT3A expression. a,** DG OE-D3A mice froze at the same rate as controls when 24h later re-exposed to the context in which fear conditioning was initially performed. Two-way ANOVA baseline vs context $F(1,16)=33.38$, $P<0.0001$. Adjusted p-value indicated on graph, n=5/group. Error bars are mean ± s.e.m. **b**, Same as in (a) but with cKO mice. Two-way ANOVA baseline vs context $F(1,38)=87.09$, $P<0.0001$; group.

**Supplementary Tables**

**Table S1** Links to Custom IGV tracks for env-DMR files

| | Custom IGV track links |
|---|---|
| run-DMRs | https://abc.med.cornell.edu/bambam_athena/stream/track/1577.bed?access_token=e024cf56c7a7a6dc364fe6106486495b |
| stress-DMRs | https://abc.med.cornell.edu/bambam_athena/stream/track/1576.bed?access_token=9c1a73663ff87df4e05895c2121d5134 |

**Table S2** Summary of changes in hyper- and hypo-methylation induced by sustained environmental exposure.

| | | run-DMR | stress-DMR |
|---|---|---|---|
| Hyper | Sites | 4091 | 2939 |
| | % total sites | 51.16 | 50.21 |
| | % shift | 27.19% | 23.13% |
| Hypo | Sites | 3905 | 2914 |
| | % total sites | 48.84 | 49.79 |
| | % shift | -27.25% | -23.48% |

**Table S3** List of differentially expressed genes following run and stress

| run-DMR genes | run-DMR genes (continued) | stress-DMR genes | stress-DMR genes (continued) |
|---|---|---|---|
| Arhgap39 | Gsk3a | Actn1 | Mecp2 |
| Asap1 | Hs3st1 | Neurl1a | Mlc1 |
| Hecw1 | Ildr2 | Orai2 | Mospd1 |
| Neurl1a | Ippk | Rcan1 | Mpzl1 |
| Npy | Kif3b | Sesn1 | Orc6 |
| Nrtn | Klk6 | Slc8a2 | Pcm1 |
| Tnfaip1 | Lars2 | Trak1 | Peli2 |
| Trak1 | Loh12cr1 | 0610030E20Rik | Pgrmc1 |
| 0610009L18Rik | Lrrc4b | 4833424O15Rik | Phf6 |
| 1110004F10Rik | Magt1 | App | Pid1 |
| 1600002K03Rik | Map2k5 | Asap1 | Pink1 |
| Actn1 | Max | Cbx7 | Pitpnb |
| Actr3b | Mcrs1 | Ccnk | Pnpla8 |
| Adcy8 | Mob2 | Cdv3 | Ppap2a |
| App | Mpzl1 | Cnih | Prepl |
| Atp13a1 | Mxi1 | Cpeb4 | Psmd10 |
| Ccdc58 | Neo1 | Crk | Ptprn |
| Cdv3 | Nrbf2 | D930007J09Rik | Puf60 |
| Clec3b | Nts | Ddb1 | Rab28 |
| Col4a3bp | Orai2 | Ddx50 | Rcsd1 |
| Cox4i2 | Orc6 | Dis3 | Rnps1 |
| D930007J09Rik | Pcdh10 | Dlst | Rpl10-ps3 |
| Dctn4 | Pcm1 | E130309D14Rik | Sgpp1 |
| Dlst | Peli2 | Fam134b | Sh2b1 |
| Efcab1 | Pid1 | Fam53c | Slc25a18 |
| Faim | Pink1 | Gkn3 | Slc25a46 |
| Fam134b | Pitpnb | Gm10291 | Sptlc1 |
| Fam53c | Pnpla8 | Gm10327 | Sult4a1 |
| Fgfrl1 | Psmd10 | Gm10358 | Syngr1 |
| Gfra2 | Puf60 | Gm10359 | Tcf12 |

| | | | |
|---|---|---|---|
| Glb1 | Pvrl4 | Gm10481 | Tmem60 |
| Gm10291 | Rab13 | Gm10566 | Tmem9b |
| Gm10293 | Rab28 | Gm12033 | Tmsb15l |
| Gm10327 | Ramp1 | Gm1673 | Trmt5 |
| Gm10358 | Rasgef1a | Gm17555 | Uba3 |
| Gm10359 | Rcsd1 | Gm2574 | Ubc |
| Gm10481 | Rnf138 | Gm266 | Ube3a |
| Gm10566 | Rps2-ps5 | Gm3200 | Usp15 |
| Gm12033 | Rps2-ps6 | Gm3222 | Usp47 |
| Gm1673 | Sgpp1 | Gm3272 | Vkorc1l1 |
| Gm17555 | Sh2b1 | Gm3839 | Zfp498 |
| Gm2574 | Slc25a18 | Gm4609 | Zfp938 |
| Gm3200 | Slc25a46 | Gm5138 | Zhx1 |
| Gm3222 | Slc36a4 | Gm5507 | |
| Gm3272 | Slc9a3r2 | Gm6139 | |
| Gm3839 | Stx6 | Gna12 | |
| Gm4609 | Sult4a1 | Gng8 | |
| Gm5138 | Syngr1 | Gpr151 | |
| Gm5507 | Tmem9b | Hs3st1 | |
| Gm5559 | Tmsb15l | Hsdl2 | |
| Gm5921 | Tpp2 | Inf2 | |
| Gm6139 | Uba3 | Kitl | |
| Gm8226 | Usp15 | Lair1 | |
| Gna12 | Usp47 | Lars2 | |
| Gnai2 | Zfp498 | Loh12cr1 | |
| Gng8 | Zfp938 | Max | |
| Gpr151 | | Mcrs1 | |

Genes that are differentially expressed that also contained DMRs are highlights in green (run-DMR) and orange (stress-DMR)

**Table S4** List of genes containing DMRs.

See included excel file

**Table S5** Summary of changes in methylation induced Dnmt3a manipulations.

| | | Clustering sites | | | Hyper | | Hypo | |
|---|---|---|---|---|---|---|---|---|
| | DMR | Average per DMR | Average length | Total number | Sites | % total sites | Sites | % total sites |
| OE-DMR | 6515 | 2.85 | 158.08 | 18576 | 17829 | 95.98 | 747 | 4.02 |
| cKO-DMR | 2920 | 3.95 | 196.58 | 11530 | 142 | 1.23 | 11388 | 98.77 |
